# Text guidance is powerful but prompt-sensitive for weakly-supervised leaf symptom segmentation

**DOI:** 10.64898/2026.07.10.737680

**Authors:** Romane Dubois, Lydia Bousset, Stéphane Jumel, Melen Leclerc, Nicolas Parisey, Alexis Joly

## Abstract

Accurate segmentation of plant disease symptoms is essential for crop monitoring and phenotyping, yet it typically requires costly pixel-level annotations. Weakly supervised semantic segmentation (WSSS) alleviates this burden using image-level labels, but its performance depends on the quality of spatial priors such as class activation maps (CAMs). We investigate whether text-guided segmentation with the Segment Anything Model 3 (SAM3) can serve as an alternative weak supervision signal. Three pseudo-mask generation strategies are compared: (i) CAMs refined with SAM or SAM3, (ii) zero-shot text-guided SAM3, and (iii) a hybrid approach combining weak spatial cues with text prompts. The resulting pseudo-masks are used to train a DeepLabV3 model. Text guidance alone matches or outperforms conventional WSSS, achieving up to 0.46 IoU without spatial supervision and 0.61 IoU on a public dataset, although performance is sensitive to text prompt formulation. The hybrid strategy improves robustness, reaching 0.50 IoU on the primary dataset and 0.58 IoU on the additional dataset while reducing prompt sensitivity. Overall, text guidance is a promising alternative to conventional weak supervision, while hybrid approaches provide a more robust solution for plant disease segmentation.

## 1. Introduction

Various pathogen agents, such as bacteria, fungi, and viruses infect plants. They can induce visible symptoms in plant organs and cause substantial yield losses. Phenotyping these symptoms accurately is essential for managing outbreaks, developing effective treatments, and identifying resistant varieties [1]. Advances in computer vision have significantly improved the phenotypic description of plant diseases, in particular through symptom segmentation tasks (e.g. [2, 3, 4, 5]).

However, although supervised approaches achieve strong performance, they rely on pixel-level annotation masks for each image. Producing these annotations is both costly and time-consuming. This difficulty is further amplified by the considerable diversity of pathosystems: symptoms vary widely in colour, texture, shape, and spatial distribution across crops and diseases, such that models trained on one context rarely transfer to another without extensive re-annotation. Yet despite this high demand for labelled data, the plant pathology community suffers from a critical shortage of openly shared datasets. The vast majority of image collections are neither publicly available nor sufficiently diverse to serve as a common benchmark for the community [6]. Altogether, these constraints strongly limit the scalability and transferability of supervised approaches to new crops, diseases, or environmental conditions, and underscore the need for methods capable of operating with reduced or no pixel-level supervision.

Several strategies have been explored to mitigate the impact of limited annotated data [7]. A first category of methods relies on data augmentation and synthetic data generation to artificially expand training sets, but these techniques remain constrained by the realism of the generated samples and do not fundamentally reduce the need for pixel-level annotations. Transfer learning and domain generalization methods offer another strategies, by leveraging knowledge acquired on large generic datasets and adapting it to plant pathology contexts. However, as noted by Wang et al. (2022) [7], these approaches often still require multiple annotated source domains to achieve robust generalization, which represents a critical bottleneck given the scarcity of labelled data. A more radical strategies consists in rethinking the supervision signal itself. Weakly supervised semantic segmentation (WSSS) has emerged in this regard as a particularly promising direction. Rather than requiring costly pixel-level masks, WSSS methods train segmentation models from coarser and cheaper forms of annotation, such as image-level labels, bounding boxes, or points which dramatically reducing annotation effort. Most approaches rely on class activation maps (CAMs) [8] to highlight discriminative regions, which are then refined into pseudo-masks, dense pixel-level labels automatically generated from weak supervision signals and used as a substitute for manual annotations during training (Figure 1). Recent works have demonstrated the potential of these methods to maintain competitive segmentation performance at a fraction of the annotation cost [9, 10], making them especially attractive for plant pathology applications.

**Figure 1:**
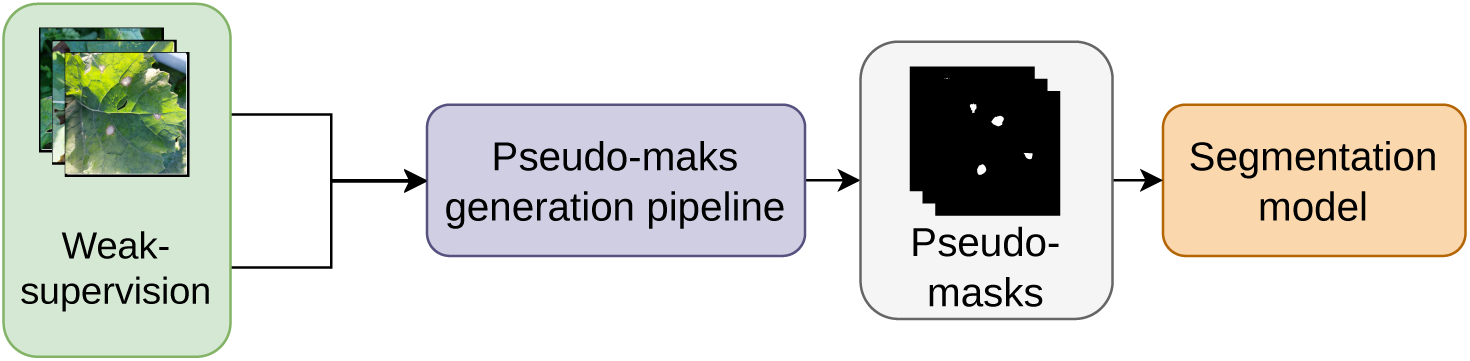
Generic weakly supervised semantic segmentation (WSSS) pipeline. A weak supervision signal (e.g. image-level label, bounding box, or text prompt) is used to generate pseudo-masks, which substitute for manual pixel-level annotations to train a segmentation model. No pixel-level annotation is required at any stage.

In the context of plant disease analysis, only a limited number of studies have explored weakly supervised strategies, despite the growing availability of image-level labeled datasets and the persistent scarcity of pixel-level annotations [11, 12, 13]. For instance, Zhou et al. (2023) [11] constructs a pixel-annotated diseased leaf dataset and implements supervised semantic segmentation (DeepLabV3+ [14]) alongside weakly supervised approaches combining Grad-CAM with ResNet-50 [15] and a few-shot pretrained U-Net[16](WSLSS), enabling pixel-level disease spot detection trained primarily on image-level labels to reduce annotation cost while evaluating cross-species generalization.

The emergence of foundation segmentation models such as the Segment Anything Model (SAM) [17] has opened new perspectives for low-annotation segmentation [18, 19]. SAM demonstrates strong zero-shot generalization capabilities and can produce high-quality object masks from simple prompts, such as points or bounding boxes [20]. These advances make SAM-based approaches especially suitable for pseudo-mask generation, where coarse object localization can be refined into accurate segmentation masks [21], a property that extends naturally to plant disease symptom segmentation. For instance, Tian et al. (2025) [22] propose UPLS, a weakly supervised pipeline for plant disease symptom segmentation that leverages PDDD-pretrained ResNet [23] to generate CAMs via contrastive learning, followed by automated SAM refinement. Building on this paradigm, newer models such as YOLOE-26[24] and SAM3[19] extend these capabilities by enabling text-guided segmentation, allowing objects of interest to be segmented based solely on semantic descriptions without explicit spatial input.

Building on these developments, this paper investigates whether text-guided weak supervision enabled by recent foundation models can serve as an alternative to, or complement, traditional spatial supervision for plant disease symptom segmentation, with the ambition of developing an approach generalizable across pathosystems. A key design principle of this work is to rely exclusively on off-the-shelf models, used without any architectural modification, so that the proposed pipelines require neither expert knowledge in deep learning nor substantial computational resources, a critical prerequisite for broad adoption across the plant pathology community.

To assess both the potential and the limitations of this approach, we first develop and evaluate three pseudo-mask generation pipelines strategies on an original dataset comprising 2540 images collected across a range of contexts (various field plots, growing seasons and environmental conditions at the time of acquisition) and covering 7 oilseed rape diseases exhibiting a variety of leaf symptoms. These pseudo-masks are used to train segmentation models evaluated along two complementary axes: segmentation quality, measured by IoU, Precision, and Recall, and symptom surface estimation, assessed via the ratio of symptomatic to total pixels and its relative Root Mean Square Error of prediction (rRMSE), directly reflecting the model’s capacity to estimate disease severity. The transferability of the three pseudo-mask generation strategies is then evaluated with the same metrics on an publicly available dataset, probing the generalization capacity of the pipelines to unseen acquisition conditions and symptom types. In addition, we examine how model performance varies depending on which annotator’s labels are used as reference, thereby assessing the extent to which evaluation outcomes are conditioned by the subjectivity inherent in manual annotation.

This paper is organized as follows. Section 2 describes the dataset and the proposed methodology. Section 3 presents the experimental results. Section 4 discusses the findings and their implications. Finally, Section 5 concludes the paper and outlines future research directions.

## 2. Materials and Methods

### 2.1. Data acquisition and manual annotation

An original dataset of 2,540 RGB images of oilseed rape leaves affected by 7 fungal and bacterial diseases (Table 1) was built to develop and evaluate the proposed pipelines, and is made publicly available [25]. These 7 diseases display a diversity of symptoms in terms of colour, texture, shape, and spatial distribution [26]. Images were acquired using smartphones under natural illumination and diverse field conditions across multiple growing seasons, from autumn 2022 to winter 2024, at the INRAE experimental station in Le Rheu (UE La Motte), France (48.1°N, 1.5°W), and in surrounding farmers’ fields. For consistency and computational efficiency, all images were center-cropped to a square format and resized to a resolution of 518 × 518 pixels.

**Table 1:** Overview of the Oilseed rape Leaf Disease Dataset, showing the number of images and pixel-level annotations per disease class.

| Pathogen | Nb. Images | Nb. Pixel-level annotations |
| --- | --- | --- |
| *Alternaria brassicae* | 328 | 93 |
| *Leptosphaeria biglobosa* | 294 | 83 |
| *Leptosphaeria maculans* | 430 | 115 |
| *Hyaloperonospora parasitica* | 401 | 73 |
| *Mycosphaerella brassicicola* | 243 | 74 |
| *Neopseudocercospora capsellae* | 351 | 81 |
| *Xanthomonas campestris* | 418 | 82 |
| Total | 2,540 | 601 |

Among these images, 580 were manually annotated at the pixel level by four annotators using CVAT[27]. To assess inter-annotator consistency detailed in Section 2.5, 21 additional images (3 per disease) were independently annotated by all four annotators. All annotations are available at https://doi.org/10.57745/OKUEDY.

### 2.2. Pseudo-mask generation strategies

Four sets of pseudo-masks were generated depending on the supervision (see Figure 2): image-level labels (two sets, see below), textual prompts (one set), and a combination of image-level labels and textual prompts (one set).

**Figure 2:**
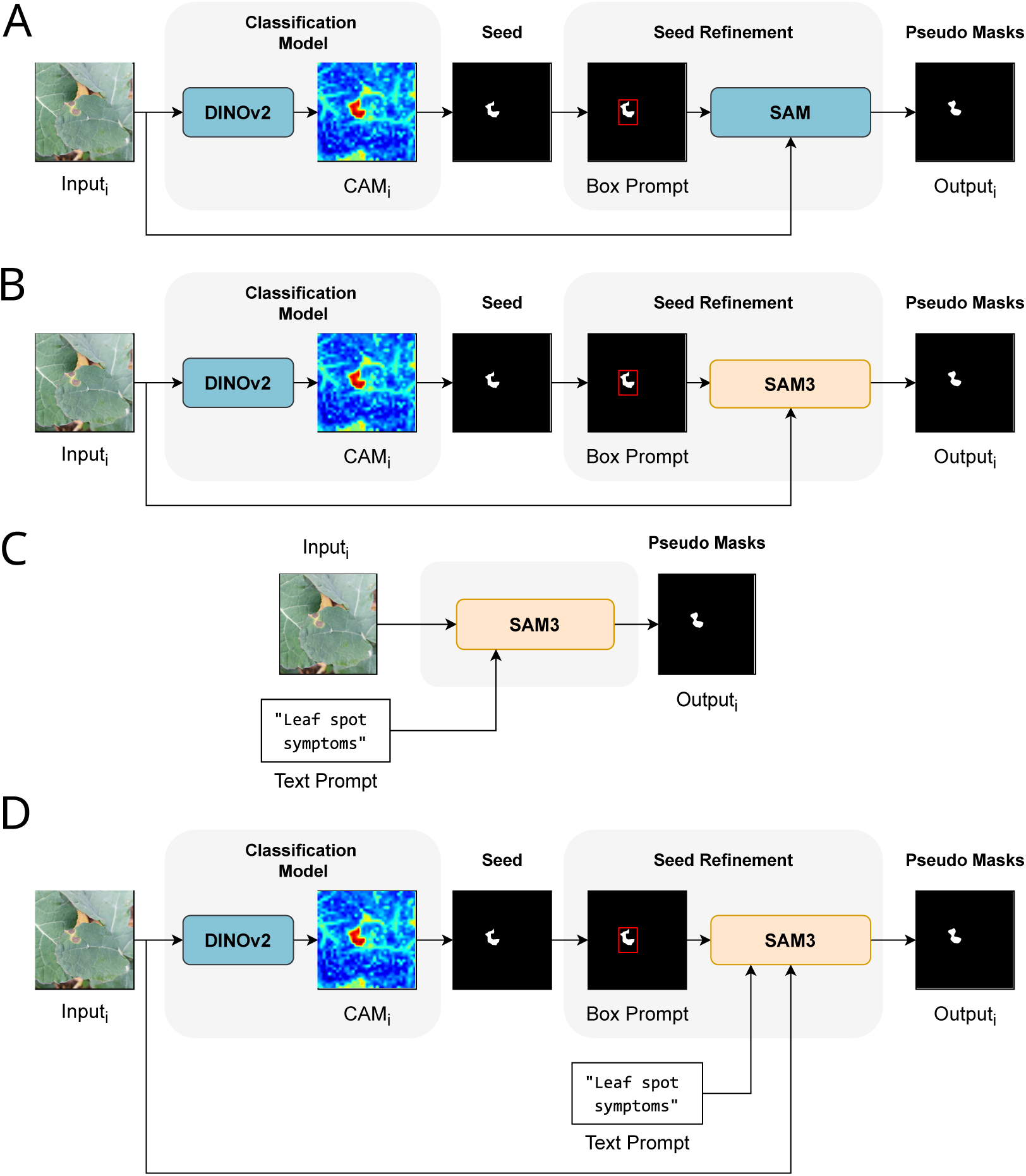
Pseudo-mask generation pipelines. (A–B) Weakly supervised approaches using DINOv2-based Class Activation Maps (CAM) as seeds, refined via SAM (A) or SAM3 (B) using a box prompt. (C) Text-guided approach using SAM3 with a text prompt only. (D) Hybrid approach combining CAM-derived box prompts and text prompts within SAM3.

#### 2.2.1. Image-level label-based pseudo-mask generation

The pipeline for the production of pseudo-masks with image-level label is divided into two steps: seed generation and seed refinement (Figure 2 A & B).

Seeds were generated using a classification model based on a DinoV2 architecture [28], a self-supervised Vision Transformer (ViT) [29] trained on a large-scale dataset of 81K images including 1.9K disease classes [30], providing rich and generalizable feature representations across a wide diversity of plant disease visual appearances.

To generate class activation maps (CAMs) from this transformer-based model, we employed Vit_CX [31], a CAM method specifically designed for Vision Transformers. Unlike gradient-based CAM methods originally developed for CNNs, Vit_CX use the attention maps of the last block of the transformer architecture, using only the patch embeddings. Preliminary experiments on our dataset showed that this approach produced more spatially precise localization of disease symptoms than GradCAM [32], likely because it better reflects the patch-based spatial structure of the ViT rather than relying on backpropagated gradients. The resulting CAMs, which produce a continuous activation map with values ranging from 0 to 1, are then binarized to generate an initial coarse segmentation. This binarization step converts the activation map into a binary mask by retaining, for each image, the top 1% most strongly activated pixels, namely those whose value exceeds the 99th percentile of that image’s own activation distribution, and setting the remaining pixels to 0, effectively selecting the most discriminative regions as belonging to the symptom class. This percentile threshold was selected based on preliminary experiments conducted on a subset of the oilseed rape dataset. The resulting binary mask constitutes the seed for the segmentation pipeline.

The binary seed masks obtained from the CAM binarization provide a coarse localization of disease symptoms. However, due to the classification objective of the underlying model, these masks often lack precise boundary delineation and may fail to capture the full spatial extent of the symptoms. To refine these seeds into more accurate segmentation masks, two segmentation models were compared: SAM [17] and SAM3 [33]. Both models are used in inference mode with box prompts: for each connected component of the binary seed mask, a bounding box prompt was automatically derived as the tightest axis-aligned rectangle enclosing the component. This bounding box is then used as a spatial prompt to guide the segmentation model toward the region of interest. The resulting segmentation output constitutes the pseudo-mask.

#### 2.2.2. Textual prompt-based pseudo-mask generation

Pseudo-masks based solely on textual prompts were generated using SAM3 [33] in inference mode, without any spatial prompt such as a bounding box (Figure 2.C). Ten different textual prompts describing leaf disease symptoms were evaluated: *lesion*, *spot*, *spots*, *leaf spot*, *leaf spots*, *leaf spot symptom*, *leaf spot symptoms*, *leaf spot disease symptom*, *leaf spot disease symptoms*, and *leaf lesion*. These prompts were intentionally restricted to generic, non-expert semantic descriptions rather than specialized pathological terminology, as SAM3 does not rely on a domain-specific vocabulary. This choice also aims to evaluate the potential transferability of text-guided segmentation to other crops, diseases, and symptom types without requiring expert-crafted prompts. The prompts were selected to cover a range of specificity levels, from generic terms such as *spot* to more descriptive phrases such as *leaf spot disease symptoms*, in order to assess the sensitivity of SAM3 to prompt formulation.

The prompt *leaf lesion* failed to generate a sufficient number of prediction masks across the dataset to constitute a large enough training set for a segmentation model. This prompt was therefore discarded, and only the remaining nine prompts were retained. Consequently, nine sets of pseudo-masks were generated and subsequently used to train nine independent segmentation models.

#### 2.2.3. Combined image-level labels and textual prompts pseudo mask generation

This hybrid approach follows the same seed generation strategy described in Section 2.2.1. For each connected component of the binary seed mask, a bounding box is automatically derived and used jointly with a textual prompt to guide SAM3 [33] toward the region of interest (Figure 2.D).

The same ten textual prompts as in Section 2.2.2 were evaluated: *lesion*, *spot*, *spots*, *leaf spot*, *leaf spots*, *leaf spot symptom*, *leaf spot symptoms*, *leaf spot disease symptom*, *leaf spot disease symptoms*, and *leaf lesion*. Unlike the purely text-based approach, the *leaf lesion* prompt produced a sufficient number of prediction masks. Consequently, all ten prompts were retained, producing ten sets of pseudo-masks, each used to train an independent segmentation model.

### 2.3. Segmentation model training

For the final segmentation task, we used a DeepLabV3 [14] architecture, a lightweight and widely adopted model for semantic segmentation, previously used in plant disease segmentation [11]. For each dataset, models were trained depending on the type of supervision used: one fully supervised model trained on manual annotations serving as an upper bound reference, and a set of weakly supervised models trained on pseudo-masks derived from seed-based prompts (2 models), text-based prompts (9 models), or a combination of both (10 models). The weakly supervised models are evaluated against the fully supervised baseline to assess the relevance of pseudo-mask-based training as an alternative to costly manual annotation.

For each dataset, a train/test split of 80/20 was applied, with images randomly assigned to each subset across the full dataset. The test set is fixed and shared across all segmentation models to ensure a fair comparison between supervision strategies. For the oilseed rape dataset, the fully supervised setting uses 464 manually annotated images for training and 116 for testing, while 2424 images are available for weakly supervised training. The effective number of training images in the text-based and hybrid settings may vary, as images for which no prediction mask was generated are excluded from training.

The batch size is set to 32, and the model parameters are optimized using the Adam optimizer over 150 epochs. The loss function is the Dice loss, commonly employed in segmentation tasks, and defined as:

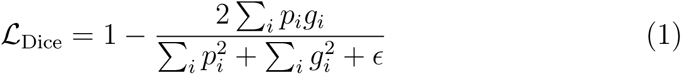

where *p_i_* and *g_i_* are respectively the predicted and ground truth values for pixel *i* and *ϵ* is a smoothing term added to ensure numerical stability.

The initial learning rate is set to *λ*_0_ = 0.1 and is adaptively reduced using a Reduce-on-Plateau strategy: if the validation loss does not decrease for 10 consecutive epochs, the learning rate is divided by a factor of 10, with a minimum learning rate of 10^−6^. An exception is made for the fully supervised model, for which a fixed learning rate of *λ* = 10^−4^ is used throughout training, along with data augmentation consisting of random horizontal and vertical flips, 90 rotations, translations, and random brightness and contrast adjustments.

These differences in training configuration reflect findings from preliminary experiments, which showed that the Reduce-on-Plateau strategy consistently improved performance for weakly supervised models, while data augmentation proved beneficial for the fully supervised model but not for weakly supervised models.

All experiments were conducted on a machine equipped with an NVIDIA RTX 4000 GPU (20 GB VRAM), an Intel Core i7-14700K CPU, and 32 GB of RAM, running Ubuntu 24.04 with Python 3.12.9, PyTorch 2.7.0, and CUDA 12.6.

### 2.4. Evaluation metrics and statistical analysis

The quality of both the pseudo-masks and the segmentation model predictions was assessed using three standard metrics commonly employed in segmentation tasks: Intersection over Union (IoU), Precision, and Recall. These metrics are computed at the pixel level, where each pixel is classified as a True Positive (TP), False Positive (FP), True Negative (TN), or False Negative (FN) based on the agreement between the predicted mask and the reference annotation.

The IoU measures the overlap between the predicted and reference masks, penalizing both over- and under-segmentation:

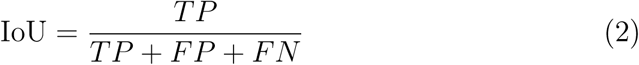

Precision measures the proportion of predicted positive pixels that are correctly classified, reflecting the ability of the model to avoid false detections:

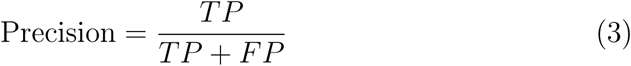

Recall measures the proportion of actual positive pixels that are correctly retrieved, reflecting the ability of the model to detect the full extent of the symptoms:

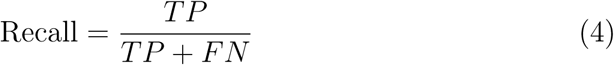

For each metric, the mean value is computed across all images of the dataset.

In addition to these metrics, the symptom coverage ratio *ρ* is computed for each image as the proportion of symptomatic pixels over the total number of pixels in the image:

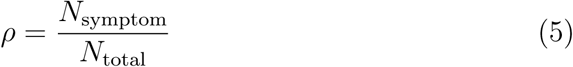

where *N*_symptom_ is the number of pixels labeled as symptomatic and *N*_total_ is the total number of pixels in the image. This ratio is computed for both the predicted masks and the reference annotations.

To evaluate the ability of each model to accurately estimate the symptomatic surface area, the relative Root Mean Square Error (rRMSE) between the predicted and annotated coverage ratios was computed across all test images:

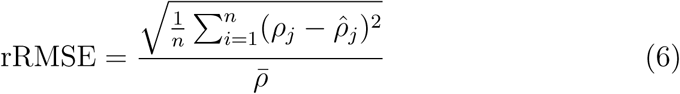

where 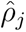 and *ρ_j_* are the predicted and annotated coverage ratios for image *j*, respectively, *n* is the total number of test images, and *ρ̄* is the mean of the annotated coverage ratios. A lower rRMSE indicates that the model produces segmentation masks whose symptomatic surface closely matches the reference annotations, relative to the average coverage ratio.

#### 2.4.1. Statistical analysis for the evaluation of segmentation performance

The IoU metric was used as the response variable in all models. Due to computational constraints, each model was trained once. Therefore, the statistical analysis captures variability across test images but does not account for training stochasticity.

To evaluate whether segmentation performance differed across the model architecture, we fitted a linear mixed-effects model on the five model configurations (FS, CAM+SAM(box), CAM+SAM3(box), CAM+SAM3(box+txt) and SAM3(txt)). For the text-based approaches CAM+SAM3(box+txt) and CAM+SAM3(txt), only the best-performing prompt, defined as the Model_Prompt combination achieving the highest mean IoU averaged across all test images, was retained for this analysis. This selection was performed on the same test set used to report final performance, rather than an independent validation split, due to the limited number of candidate prompts (9–10) and the already constrained size of the test set. The practical implication of this limitation for real-world deployment is discussed in Section 5. The model is defined as follows:

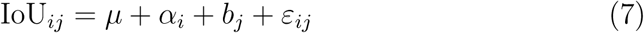

where *µ* is the overall intercept, *α_i_* is the fixed effect of model *i*, *b_j_* ∼ 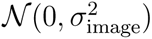 is the random effect of image *j*, and *ε_ij_* ∼ N (0*, σ*^2^) is the residual error. The random effect of image accounts for the repeated evaluation of the same test images across models.

For the text-based and hybrid models, a second linear mixed-effects model was fitted to jointly assess the influence of the model and the prompt formulation on IoU:

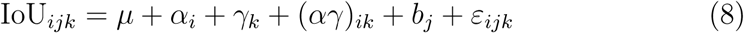

where *α_i_* is the fixed effect of model *i*, *γ_k_* is the fixed effect of prompt *k*, (*αγ*)*_ik_* is the interaction term between model and prompt, 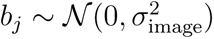 is the random effect of image *j*, and *ε_ijk_* ∼ N (0*, σ*^2^) is the residual error. The interaction term allows us to assess whether the effect of prompt formulation differs between the text-based and hybrid models.

### 2.5. Annotation consistency and its influence on segmentation evaluation

Ground-truth annotations are inherently conditioned by the annotator’s perception, introducing a degree of subjectivity that directly affects model evaluation. To assess the extent to which segmentation performance metrics depend on the choice of reference annotator, the five pipelines (FS, CAM+SAM, CAM+SAM3, SAM3(txt), CAM+SAM3(box+txt)) were evaluated independently against each annotator’s masks, using IoU as the evaluation metric. To quantify annotation variability, 21 images (3 per disease class, selected to cover different levels of disease severity) were independently annotated by all four annotators. Two complementary measures were computed. Pairwise IoU was calculated between each pair of annotators and averaged across all pairs, reflecting the typical degree of overlap between any two individual annotations. Consensus IoU was computed using the intersection of all four annotators’ masks as the reference, retaining only pixels unanimously labeled as symptomatic. By construction more conservative than pairwise IoU, this measure captures how much each individual annotation deviates from the region of full consensus, directly reflecting the extent of annotation variability across annotators.

### 2.6. Generalization and comparison with the literature

To evaluate the generalization and consistency of the developed methods with the literature, the pipelines were also applied to a public dataset used in other studies [22, 11]. The apple Leaf Disease Dataset (available at https://data.mendeley.com/datasets/tsfxgsp3z6) comprises 865 diseased apple leaf images spanning 5 disease classes (Table 2). Part of this dataset consists of standardized images from PlantVillage dataset (available at https://data.mendeley.com/datasets/tywbtsjrjv/1), while the remaining two disease classes contain field-collected images.

**Table 2:** Overview of the Apple Leaf Disease Dataset, showing the number of images per disease class. All images have pixel-level annotations.

| Disease | Image Source | Nb. Images |
| --- | --- | --- |
| Alternaria blotch | PlantVillage | 136 |
| Apple scab | PlantVillage | 156 |
| Black rot | PlantVillage | 163 |
| Cedar apple rust | Field | 98 |
| Rust | Field | 312 |
| Total |  | 865 |

Following the same experimental protocol as for the oilseed rape dataset, segmentation models were trained on each set of pseudo-masks generated by the different pipelines, using an 80/20 train/test split. The same hyperparameters were used throughout, both for the CAM binarization threshold and for the training of the segmentation models, as described in Sections 2.3 and 2.4. A fully supervised model trained on the available pixel-level annotations was also included as an upper bound reference. Segmentation performance was assessed using IoU, Precision, and Recall, allowing direct comparison across supervision strategies and with methods from the literature.

Comparisons with other methods reported in the literature were restricted to descriptive analyses, as only aggregated mean performance values are available from published studies, making impossible any inferential statistical comparison.

## 3. Results

### 3.1. Segmentation performance of each model as a function of the supervision used for training

Table 3 presents the comparison of segmentation performance using the DeepLabV3 architecture under the different supervision strategies: fully supervised, weakly supervised using image levels labels, text prompts and an hybrid approach combining both.

**Table 3:** Comparison of segmentation models (DeepLabV3+) under different supervision strategies, evaluated on the oilseed rape dataset. Prompt variability is reported for both text-guided strategies as the IoU range (min, mean, max). Segmentation performance is assessed through IoU, Precision, and Recall, while symptomatic surface estimation accuracy is evaluated using rRMSE.

| Supervision | Pseudo-Mask Pipeline | #Valid Prompts | Prompt Variability (IoU Range) | Segmentation performance |  |  | Surface estimation |
| --- | --- | --- | --- | --- | --- | --- | --- |
|  |  |  |  | ↑ IoU | ↑ Precision | ↑ Recall | ↓ rRMSE |
| Fully Supervised | – | – | – | 0.60 | 0.71 | 0.70 | 0.4808 |
| Weakly Supervised (image-level labels) | CAM + SAM(box) | – | – | 0.42 | 0.80 | 0.51 | 1.2043 |
|  | CAM + SAM3(box) | – | – | 0.44 | 0.60 | 0.70 | 0.9479 |
| Weakly Supervised (text prompt) | SAM3(txt) | 9/10 | max | 0.46 | 0.65 | 0.64 | 0.7693 |
|  |  |  | mean | 0.36 | 0.63 | 0.47 | 1.2135 |
|  |  |  | min | 0.19 | 0.48 | 0.25 | 1.1036 |
| Weakly Supervised (image-level labels + text prompt) | CAM + SAM3(box+txt) | 10/10 | max | **0.50** | **0.69** | **0.68** | **0.7739** |
|  |  |  | mean | 0.47 | 0.68 | 0.66 | 0.9232 |
|  |  |  | min | 0.44 | 0.60 | 0.69 | 1.1585 |

The fully supervised (FS) model established the baseline upperbound performance with IoU = 0.60, precision = 0.71, recall = 0.70. The weakly supervised method based on image-level labels doesn’t show signicant difference of IoU depending on the SAM version employed (p > 0.05). However CAM+SAM3(box) gets a better balanced precision (0.60) and recall (0.70), than CAM+SAM(box). Incorporating text prompts further improves segmentation performance, with both text-guided strategies achieving significantly higher IoU scores than CAM-based approaches when using the best prompt configuration (p < 0.05). Among these, the hybrid approach CAM+SAM3(box+txt) achieves the best overall weakly supervised performance (IoU = 0.50, precision = 0.69, recall = 0.68 with the prompt ’spots’), significantly outperforming the text-only SAM3(txt) (p < 0.05). Additionally, the text-only approach exhibits substantially higher sensitivity to prompt choice (IoU range: 0.19–0.46, mean = 0.36), whereas the hybrid approach demonstrates greater robustness to prompt variability (IoU range: 0.44–0.50, mean = 0.47), suggesting that combining image-level labels with text prompts not only boosts performance but also stabilizes it across different prompt formulations (see Appendix A.2 for detailed per-prompt results).

### 3.2. Symptom surface estimation performance of each model as a function of the supervision used for training

Beyond segmentation performance, we evaluate the capacity of each pipeline to estimate symptomatic surface, using the rRMSE as a measure of estimation error relative to the observed range (Table 3). The fully supervised model achieved the lowest estimation error (rRMSE = 0.48). CAM+SAM(box) showed the highest estimation error (rRMSE = 1.20), while CAM+SAM3(box) improved on this (rRMSE = 0.95). Text-guided approaches yielded intermediate values, with both the best-prompt SAM3(txt) and the best-prompt CAM+SAM3(box+txt) reaching rRMSE = 0.77. All models systematically underestimate symptomatic surface(see Appendix D.2).

### 3.3. Annotation consistency and its influence on segmentation evaluation

We computed the IoU of each pipeline against each annotator’s masks independently across the 21 images annoted (Table 4). Model rankings remain stable across annotators, and absolute IoU values vary by up to 0.05 points depending on the reference annotator. To investigate the source of this variability, we assessed inter-annotator consistency among each annotators using consensus IoU and pairwise agreement. Consensus IoU averaged 0.59, while pairwise IoU averaged 0.74 (Table 5). Figure 3 illustrates two key challenges in manual annotation: detecting small and subtle lesions leads to inter-annotator variability in symptom count (Example A), while precise boundary delimitation remains inconsistent even for larger, more visible symptoms (Example B).

**Figure 3:**
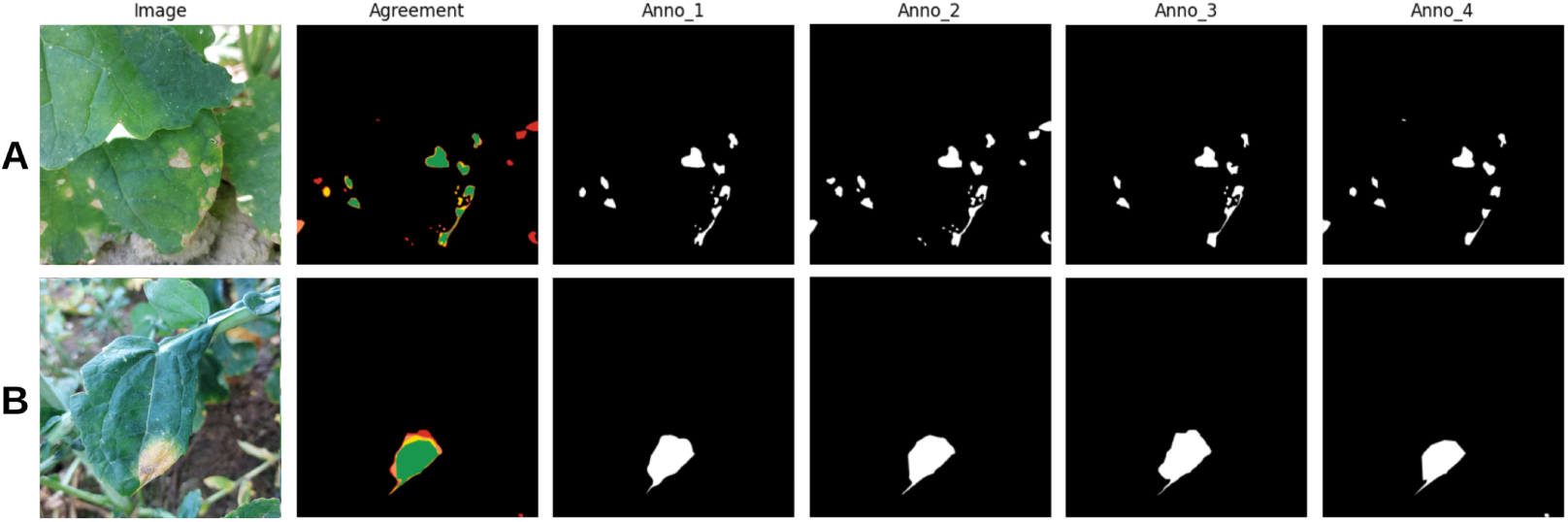
Qualitative comparison of the annotation step by different annotators for two example images A and B. On the agreement mask, each color corresponds to the number of annotators who agreed to classify the pixel as the symptom class (green = 4, yellow = 3, orange = 2, red = 1, black = 0)

**Table 4:** IoU of each pipeline evaluated against each annotator’s masks independently.

| Pipeline | Anno1 | Anno2 | Anno3 | Anno4 |
| --- | --- | --- | --- | --- |
| FS | 0.61 | 0.59 | 0.59 | 0.56 |
| CAM+SAM | 0.42 | 0.41 | 0.41 | 0.39 |
| CAM+SAM3 | 0.47 | 0.46 | 0.45 | 0.43 |
| SAM3(txt) | 0.55 | 0.55 | 0.55 | 0.53 |
| CAM+SAM3(box+txt) | 0.53 | 0.52 | 0.52 | 0.50 |

**Table 5:**
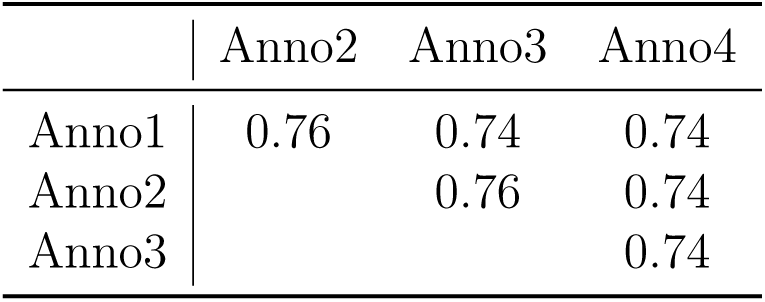
Inter-annotator agreement matrix (IoU). Diagonal omitted.

**Table 6:** Comparison of segmentation models (DeepLabV3 architecture) under different supervision strategies on the apple leaf dataset (publicly available). The values for WSLSS-400 [11], ResNet-CAM [11], and UPLS [22] are taken from Table 2 of [22]. Values for WSLSS-400, ResNet-CAM, and UPLS are taken directly from [22] and were obtained with different train/test splits, backbone architectures, and training procedures than those used in this study. These comparisons are therefore descriptive only; no inferential statistical test was performed across studies. Moreover although presented as “unsupervised,” UPLS fundamentally relies on the initial supervision of 400k PDDD images, and can therefore be more accurately categorized as weakly supervised.

| Supervision | Pseudo-Masks pipeline | #Valid Prompts | Prompt variability (IoU Range) | ↑ IoU | ↑ Precision | ↑ Recall |
| --- | --- | --- | --- | --- | --- | --- |
| Fully Supervised | – | – | – | 0.72 | 0.84 | 0.83 |
| Semi Supervised | WSLSS-400 [11] | – | – | 0.43 | 0.74 | 0.58 |
| Weakly Supervised (image-level labels) | ResNet-CAM [11] | – | – | 0.20 | 0.30 | 0.50 |
|  | UPLS [22] | – | – | 0.48 | 0.74 | 0.60 |
|  | CAM+SAM(box) (ours) | – | – | 0.28 | 0.58 | 0.33 |
|  | CAM+SAM3(box) (ours) | – | – | 0.33 | 0.53 | 0.41 |
| Weakly Supervised (text prompts) | SAM3(txt) (ours) | 9/10 | max | 0.61 | 0.87 | 0.67 |
|  |  |  | mean | 0.48 | 0.81 | 0.54 |
|  |  |  | min | 0.33 | 0.80 | 0.36 |
| Weakly Supervised (image-level labels + text prompts) | CAM+SAM3(box+txt) (ours) | 10/10 | max | 0.58 | 0.83 | 0.67 |
|  |  |  | mean | 0.45 | 0.73 | 0.58 |
|  |  |  | min | 0.26 | 0.56 | 0.33 |

### 3.4. Generalization and comparison with the literature

To assess the generalization capacity of the proposed pipelines beyond the oilseed rape dataset, we evaluate them on an independent publicly available apple disease dataset and compare their performance against existing weakly and semi supervised methods from the literature ([11, 22]). This cross-domain evaluation allows us to probe whether pipelines developed on one crop can transfer to a different host species and symptom types without any retraining or fine-tuning.

The fully supervised model again achieved the highest performance (IoU = 0.72). Unlike the oilseed rape dataset, CAM-based only strategies differed significantly from each other (p < 0.05), with CAM+SAM3(box) slightly outperforming CAM+SAM(box) (IoU = 0.33 vs. 0.28). Again incorporating text prompts improves segmentation performance, with both text-guided strategies achieving significantly higher IoU scores than CAM-based approaches when using the best prompt configuration (p < 0.05). However, unlike the oilseed rape dataset, the text-only approach SAM3(txt) and the hybrid CAM+SAM3(box+txt) achieve comparable best-prompt performance (IoU = 0.61 vs. 0.58, respectively), with no statistically significant difference between them (p > 0.05). Both models showed comparable prompt sensitivity (IoU range: 0.26-0.58 for CAM+SAM3(box+txt) vs. 0.33-0.61 for SAM3(txt)), suggesting that the bounding box did not stabilize performance across prompts on this dataset.

Compared to some weakly and semi supervised from the literature, our CAM-based only approaches remained below UPLS (IoU = 0.48), with CAM+SAM(box) (IoU = 0.28) and CAM+SAM3(box)(IoU = 0.33). However, our text-guided methods achieved competitive results: SAM3(txt) reached a best-prompt IoU = 0.61 and CAM+SAM3(box+txt) a best-prompt IoU = 0.58, both substantially outperforming UPLS and approaching the fully supervised baseline (IoU = 0.72).

## 4. Discussion

This study investigates whether text-guided segmentation can serve as an efficient alternative to conventional weakly supervised approaches based on image-level labels for plant disease symptom segmentation. Three pseudo-mask generation strategies were developed on an oilseed rape leaf disease dataset and their generalization capacity assessed on an additional publicly available apple leaf disease dataset, revealing distinct performance profiles across image domains and supervision types.

### 4.1. Fully supervised performance and annotation reliability

In both datasets, the fully supervised baseline achieved the highest segmentation performance (IoU = 0.60 in oilseed rape, IoU = 0.72 in apple), confirming that pixel-level annotations remain the standard for training segmentation models [9, 10]. However, this performance remains dependent on annotation quality, which is itself annotator-dependent. This variability arises from two sources: the difficulty of detecting subtle lesions, leading to inconsistencies in symptom count, and the challenge of delineating symptom boundaries (Figure 3). Consensus IoU among four annotators averaged 0.59, while pairwise IoU averaged 0.74. This human plateau of 0.59–0.74 demonstrates that manual annotation introduces an irreducible degree of subjectivity, which directly conditions the reliability of ground-truth annotations and, consequently, the performance of fully supervised models. However, it should be noted that with only 21 annotated images, the statistical power remains limited, preventing a deeper exploration of both the variation in model performance as a function of the reference annotator and the variability in annotations across annotators. A larger multi-annotator dataset would be required to draw robust conclusions on these aspects. A larger multi-annotator dataset would allow this long-standing and well-documented issue to be addressed more rigorously. Broadly, annotator variability can be treated either as noise obscuring a single hidden ground truth, to be estimated, or as a signal to be predicted in its own right. The former is illustrated by statistical consensus methods such as the Simultaneous Truth and Performance Level Estimation (STAPLE) algorithm [34], which models the unknown ground truth as a latent variable and jointly estimates each annotator’s sensitivity and specificity through an expectation-maximization framework, rather than relying on a single reference annotator or a simple majority vote. The latter is exemplified by probabilistic generative architectures [35], which directly model the distribution of plausible segmentations induced by multiple annotators.

### 4.2. Prompt sensitivity as an inherent limitation of text-guided segmentation

The sensitivity to text prompts formulation is an inherent limitation of SAM3 : noun phrases are encoded as raw sub-word sequences without linguistic normalization [33], so semantically close formulations such as *spot* and *spots* produce distinct representations inducing different segmentation behaviors. On the oilseed rape dataset, SAM3(txt) exhibited substantial variability (IoU range: 0.19–0.46, mean = 0.36), while the hybrid CAM+SAM3(box+txt) partially attenuated this sensitivity (IoU range: 0.44-0.50, mean = 0.47). A concrete illustration is the *leaf lesion* prompt, which failed to generate sufficient number of masks in the text-only configuration but succeeded in the hybrid one, where the bounding box compensated for its weak discriminative power. In a deployment scenario where the optimal prompt for a new pathosystem is unknown, the hybrid approach therefore acts as a safety net, reducing the risk of prompt-induced failure without requiring any additional annotation. However, this stabilizing effect is conditioned on the quality of the CAM-derived bounding boxes: as shown on the apple dataset, when CAM quality degrades, the hybrid approach loses this advantage and no longer outperforms the text-only configuration, making SAM3(txt) with careful prompt selection the more robust strategy in cross-domain settings.

### 4.3. Generalization capacity of text-guided versus image-level label approaches

CAM-based approaches showed limited generalization on the apple dataset (CAM+SAM(box): IoU = 0.28, CAM+SAM3(box): IoU = 0.33), likely because the PlantVillage standardized images it contains are underrepresented in large-scale plant disease datasets, resulting in less informative CAMs. This degradation may be compounded by the fact that the CAM binarization threshold (0.99) was calibrated on the oilseed rape dataset and reused without adaptation on apple; a threshold optimized for one image domain’s typical activation distribution may be poorly suited to another. Any degradation in CAM quality propagates downstream to the pseudo-masks, highlighting a fundamental dependency of CAM-based pipelines on the representativeness of the classification backbone’s training domain. A low-cost avenue to partially mitigate this dependency, requiring no architectural change or additional annotation, lies in the CAM binarization threshold, held fixed here: an adaptive per-image threshold could better accommodate variable lesion sizes and contrasts, and represents a logical next step before more costly interventions [13]. In contrast, text-guided approaches generalized substantially better, with SAM3(txt) reaching IoU = 0.61 on the apple dataset, surpassing all weakly supervised baselines and approaching the fully supervised upper limit (0.72). This advantage stems from the broad visual and linguistic knowledge encoded in SAM3 during large-scale pretraining, which does not rely on domain-specific feature representations and is therefore less sensitive to distributional shifts between datasets. This result should nonetheless be interpreted as a best-prompt performance, underlining that prompt selection remains the key operational lever in annotation-free deployment. Beyond prompt optimization, all experiments were conducted using DeepLabV3+ as the segmentation backbone; replacing it with more recent transformer-based architectures such as SegFormer [36] could potentially improve segmentation performance across all supervision strategies, and represents a natural direction for future work.

### 4.4. Systemic underestimation of disease severity

All pipelines systematically underestimated symptomatic surface, following the same ranking as segmentation performance. This reflects a structural property of CAM-based training: activations concentrate on the most discriminative regions, causing pseudo-masks to undersample the full lesion extent [8, 13]. Any pipeline deployed for severity scoring would therefore require an explicit post-hoc calibration step, for instance fitting a correction factor or regression model that maps the predicted symptomatic surface to the true surface on a held-out annotated subset, in order to compensate for this systematic underestimation before the score is used for diagnosis. Beyond calibration, converting symptomatic surface into a true agronomic severity score requires normalizing by total leaf surface. Interestingly, [5] report that this ratio does not suffer from the same underestimation bias.

## 5. Conclusion

This study evaluated three pseudo-mask generation strategies for weakly supervised plant disease symptom segmentation, comparing CAM-based approaches with text-guided and hybrid pipelines built on SAM and SAM3. Text-guided segmentation achieved competitive performance without any pixel-level annotations, reaching IoU = 0.61 on the apple dataset and outperforming all weakly and semi-supervised baselines, while the hybrid approach reached IoU = 0.50 on the oilseed rape dataset with greater robustness. However, prompt formulation remains a critical unresolved hyperparameter. In this study, the best prompt was identified directly on the test set (Section 2.4), which optimistically biases the reported performance. In a real deployment setting where ground-truth masks are unavailable, prompt selection would instead require a small annotated validation set, partially undermining the annotation-free premise of the approach. All models systematically underestimated disease severity, requiring explicit calibration before use in quantitative phenotyping pipelines. This study establishes a foundation for integrating text-guided segmentation into plant disease monitoring workflows, highlighting that closing the gap with fully supervised approaches depends less on architectural improvements than on resolving the practical challenges of prompt selection.

## Acknowledgments

The authors thank Théo Fabien for managing the CVAT annotation server. This work was supported by the French National Research Agency (ANR) [Pl@ntAgroEco - ANR-22-PEAE-0009].The funder had no role in the design of the study; in the collection, analysis, or interpretation of data; in the writing of the manuscript; or in the decision to publish the results.

## Declaration of generative AI use

During the preparation of this work, the authors used generative artificial intelligence tools to improve the clarity and language of the text. The authors reviewed and edited the output as needed and take full responsibility for the content of the published article.

