## Supplementary_Materials for "Text guidance is powerful but prompt-sensitive for weakly-supervised leaf symptom segmentation"

### A Segmentation performance of each model on Oilseed rape dataset - Statistical analysis details

#### A.1 Effect of model on segmentation performance across model configurations

Table 1: Type III Analysis of Variance table for the linear mixed-effects model assessing the effect of Model (FS, SAM(box), SAM3(box), SAM3(box+txt), SAM3(txt)) on segmentation performance (IoU).

| | Sum Sq | Mean Sq | NumDF | DenDF | F value | Pvalue |
|---|---|---|---|---|---|---|
| Model | 2.42 | 0.61 | 4.00 | 460.00 | 57.07 | <0.0001 |

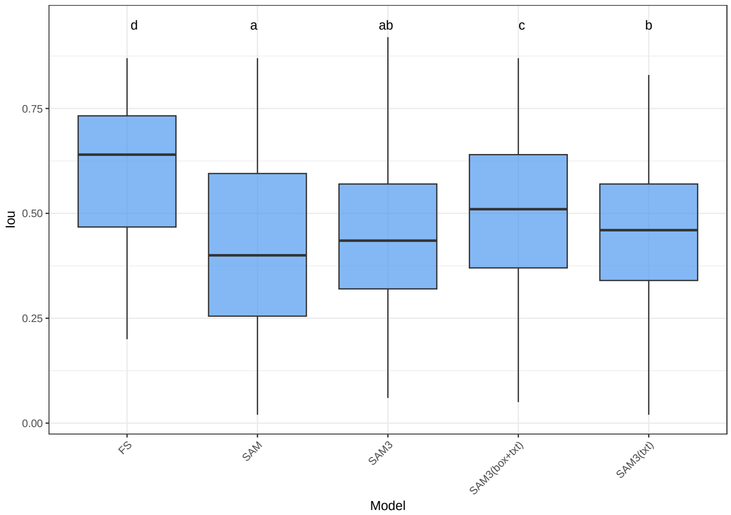


Figure 1: Segmentation performance (IoU) of models FS, SAM, SAM3, SAM3(box+txt) and SAM3(txt). Boxplots show the distribution of IoU values, letters above boxes indicate homogeneous groups with no significant difference from post-hoc pairwise comparisons.

#### A.2 Effect of prompt on segmentation performance across Model_Prompt combinations

Table 2: Type III Analysis of Variance table for the linear mixed-effects model assessing the effects of Model (SAM3(txt), SAM3(box+txt)), Prompt, and their interaction on segmentation performance (IoU).

| | Sum Sq | Mean Sq | NumDF | DenDF | F value | Pvalue |
|---|---|---|---|---|---|---|
| Model | 6.60 | 6.60 | 1.00 | 2070.00 | 440.01 | <0.0001 |
| Prompt | 5.54 | 0.62 | 9.00 | 2070.00 | 41.04 | <0.0001 |
| Model:Prompt | 4.64 | 0.58 | 8.00 | 2070.00 | 38.62 | <0.0001 |

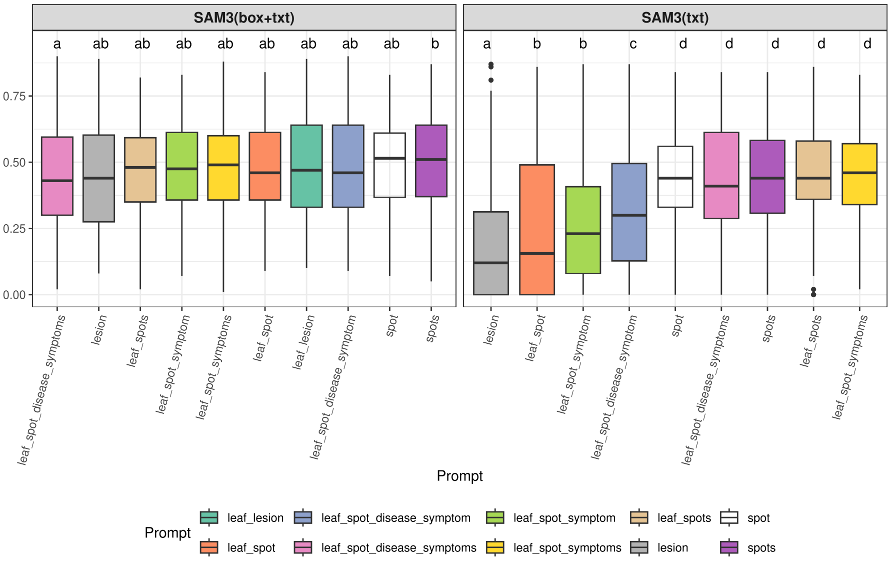


Figure 2: Segmentation performance (IoU) of models SAM3(box+txt) and SAM3(txt) depending of text prompts. Boxplots show the distribution of IoU values, letters above boxes indicate homogeneous groups with no significant difference from post-hoc pairwise comparisons.

#### B Segmentation performance of each model on Apple leaf Dataset - Statistical analysis details

##### B.1 Effect of model on segmentation performance across model configurations on Apple dataset

| | Sum Sq | Mean Sq | NumDF | DenDF | F value | Pvalue |
|---|---|---|---|---|---|---|
| Model | 25.06 | 6.27 | 4.00 | 688.00 | 173.06 | 0.0000 |

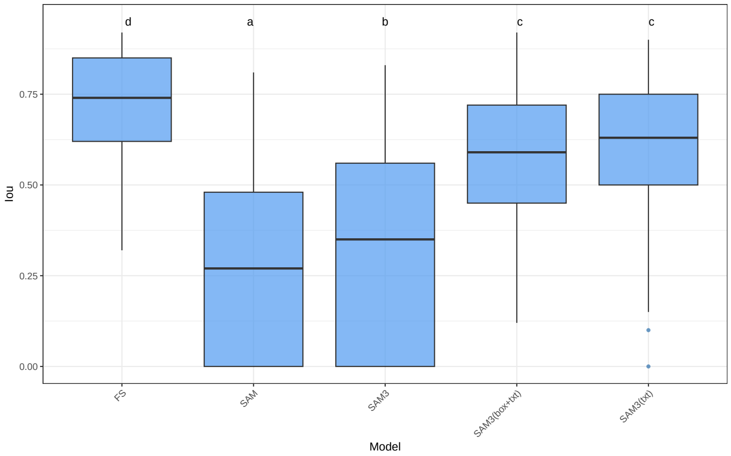


Figure 3: Segmentation performance (IoU) of models FS, SAM, SAM3, SAM3(box+txt) and SAM3(txt). Boxplots show the distribution of IoU values, letters above boxes indicate homogeneous groups with no significant difference from post-hoc pairwise comparisons.

##### B.2 Effect of prompt on segmentation performance across Model_Prompt combinations

| | Sum Sq | Mean Sq | NumDF | DenDF | F value | Pvalue |
|---|---|---|---|---|---|---|
| Model | 0.21 | 0.21 | 1.00 | 3096.00 | 6.19 | 0.0129 |
| Prompt | 33.10 | 3.68 | 9.00 | 3096.00 | 111.10 | 0.0000 |
| Model:Prompt | 5.33 | 0.67 | 8.00 | 3096.00 | 20.12 | 0.0000 |

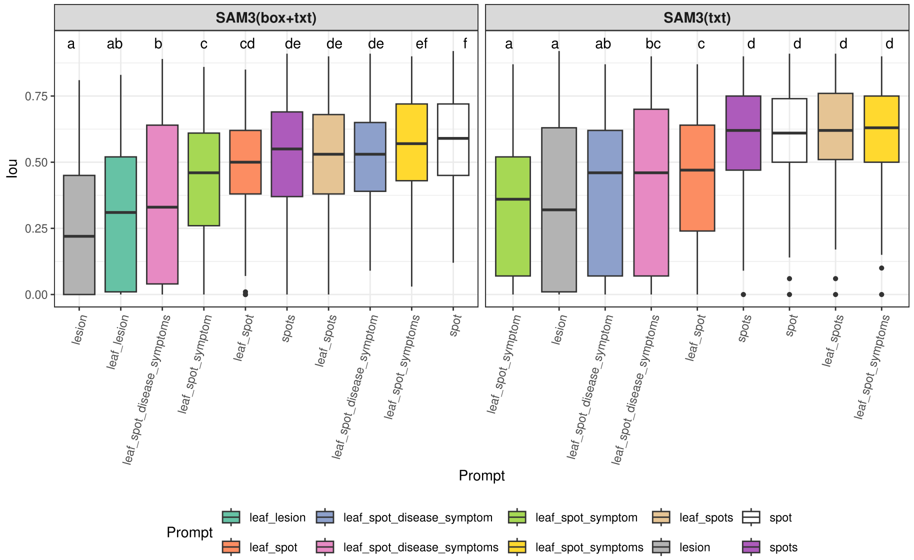


Figure 4: Segmentation performance (IoU) of models SAM3(box+txt) and SAM3(txt) depending of text prompts. Boxplots show the distribution of IoU values, letters above boxes indicate homogeneous groups with no significant difference from post-hoc pairwise comparisons.

### C Models segmentation performance (IoU) per oilseed rape leaf disease

Segmentation performance was evaluated across seven oilseed rape leaf diseases for five model configurations: one fully supervised baseline (FS) and four weakly supervised approaches (SAM, SAM3, SAM3(box+txt) and SAM3(txt)). Among these, SAM and SAM3 rely solely on geometric prompts (bounding boxes), while SAM3(box+txt) and SAM3(txt) additionally incorporate textual prompts. The two text-based approaches were used to train multiple models: 10 models were trained with SAM3(box+txt) and 9 with SAM3(txt), as one prompt failed to produce a viable model, resulting in a total of 19 Model_Prompt combinations. Two complementary analyses were conducted: the first compared the five model configurations using the best-performing prompt for each text-based approach, defined as the Model_Prompt combination achieving the highest mean IoU averaged across all test images and disease types; the second extended this analysis to all 19 Model_Prompt combinations to investigate whether the sensitivity to disease type differed according to prompt formulation.

#### C.1 Effect of disease across model configurations

To evaluate whether segmentation performance differed across oilseed rape leaf diseases and whether this effect depended on the model architecture, we fitted a linear mixed-effects model on the five model configurations (FS, SAM, SAM3, SAM3(box+txt) and SAM3(txt)). For the text-based approaches SAM3(box+txt) and SAM3(txt), only the best-performing prompt, defined as the Model_Prompt combination achieving the highest mean IoU averaged across all test images and disease types, was retained for this analysis. The model is defined as follows:

$$\mathrm{IoU}_{ijk} = \mu + \alpha_i + \beta_j + (\alpha\beta)_{ij} + b_k + \varepsilon_{ijk} \tag{1}$$

where $\mu$ is the overall intercept, $\alpha_i$ is the fixed effect of model $i$ (FS, SAM, SAM3, SAM3(box+txt), SAM3(txt)), $\beta_j$ is the fixed effect of disease $j$, $(\alpha\beta)_{ij}$ is the interaction between model $i$ and disease $j$, $b_k \sim \mathcal{N}(0, \sigma_b^2)$ is the random effect of image $k$, and $\varepsilon_{ijk} \sim \mathcal{N}(0, \sigma^2)$ is the residual error.

Table 3: Type III Analysis of Variance table for the linear mixed-effects model assessing the effects of Model (FS, SAM, SAM3, SAM3(box+txt), SAM3(txt)), disease, and their interaction on segmentation performance (IoU).

| | Sum Sq | Mean Sq | NumDF | DenDF | F value | Pvalue |
|---|---|---|---|---|---|---|
| Model | 2.37 | 0.59 | 4.00 | 436.00 | 61.22 | 0.0000 |
| Disease | 0.35 | 0.06 | 6.00 | 109.00 | 6.10 | 0.0000 |
| Model:Disease | 0.66 | 0.03 | 24.00 | 436.00 | 2.84 | 0.0000 |

Results (Table 3) revealed significant main effects of both Model ($p < 0.05$) and Disease ($p < 0.05$), indicating that segmentation performance varied substantially across model architectures and disease types. The interaction term Model:Disease was also significant ($p < 0.05$), indicating that the effect of disease on IoU was not uniform across model architectures, meaning that certain models were more sensitive to disease type than others. Pairwise post-hoc comparisons of IoU across diseases and models are presented in Figure 5.

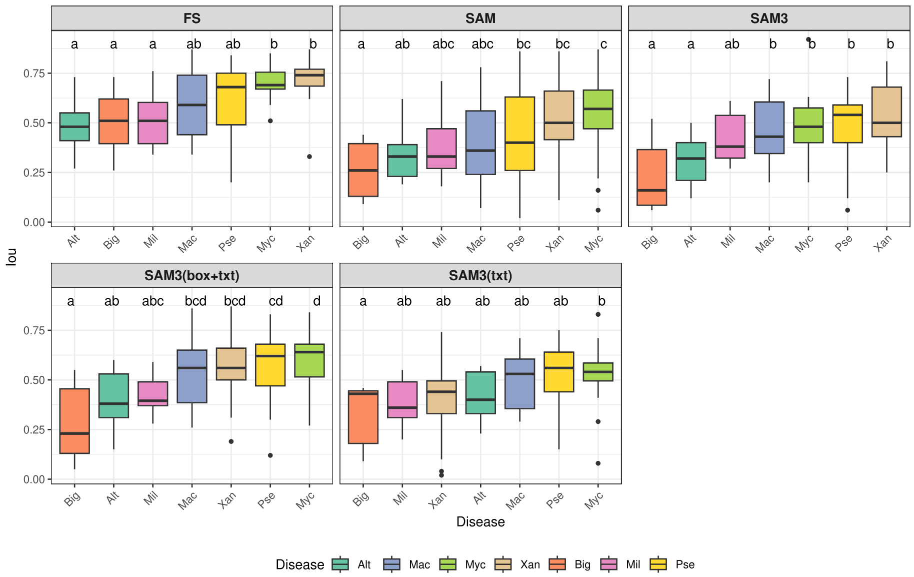


Figure 5: Segmentation performance (IoU) of models FS, SAM, SAM3, SAM3(box+txt) and SAM3(txt) per oilseed rape leaf disease. Boxplots show the distribution of IoU values, letters above boxes indicate homogeneous groups with no significant difference from post-hoc pairwise comparisons.

#### C.2 Effect of disease across Model_Prompt combinations

To investigate whether the sensitivity to disease type differed across the 19 Model_Prompt combinations derived from the text-based approaches (10 SAM3(box+txt) and 9 SAM3(txt)), a second linear mixed-effects model was fitted with `Model_Prompt`, disease, and their interaction as fixed effects. This analysis specifically captures the variability introduced by prompt formulation within each text-based architecture, which cannot be assessed from the first model alone. The model is defined as follows:

$$\mathrm{IoU}_{ijk} = \mu + \gamma_i + \beta_j + (\gamma\beta)_{ij} + b_k + \varepsilon_{ijk} \tag{2}$$

where $\mu$ is the overall intercept, $\gamma_i$ is the fixed effect of Model_Prompt combination $i$ ($i = 1, \ldots, 19$), $\beta_j$ is the fixed effect of disease $j$, $(\gamma\beta)_{ij}$ is the interaction term between Model_Prompt $i$ and disease $j$, $b_k \sim \mathcal{N}(0, \sigma_b^2)$ is the random effect of image $k$, and $\varepsilon_{ijk} \sim \mathcal{N}(0, \sigma^2)$ is the residual error. The interaction term $(\gamma\beta)_{ij}$ specifically allows us to test whether the ranking of diseases in terms of IoU is consistent across all Model_Prompt combinations, or whether certain combinations are more sensitive to disease type than others.

Table 4: Type III Analysis of Variance table for the linear mixed-effects model assessing the effects of Model_Prompt ($i = 1, \ldots, 19$), disease, and their interaction on segmentation performance (IoU).

| | Sum Sq | Mean Sq | NumDF | DenDF | F value | Pvalue |
|---|---|---|---|---|---|---|
| Model_Prompt | 15.92 | 0.88 | 18.00 | 1962.00 | 72.72 | 0.0000 |
| Disease | 0.49 | 0.08 | 6.00 | 109.00 | 6.74 | 0.0000 |
| Model_Prompt:Disease | 7.19 | 0.07 | 108.00 | 1962.00 | 5.48 | 0.0000 |

Consistent with the first model, all three terms were highly significant (Table 4). The significant main effects of Model_Prompt ($p < 0.05$) and Disease ($p < 0.05$) confirm that both prompt formulation and disease type independently contributed to variation in segmentation performance. The significant interaction term ($p < 0.05$) further indicates that the effect of disease on IoU was not consistent across Model_Prompt combinations, meaning that certain prompt strategies led to models more sensitive to disease type than others. Post-hoc pairwise comparisons for each Model_Prompt combination are visualised in Figure 6.

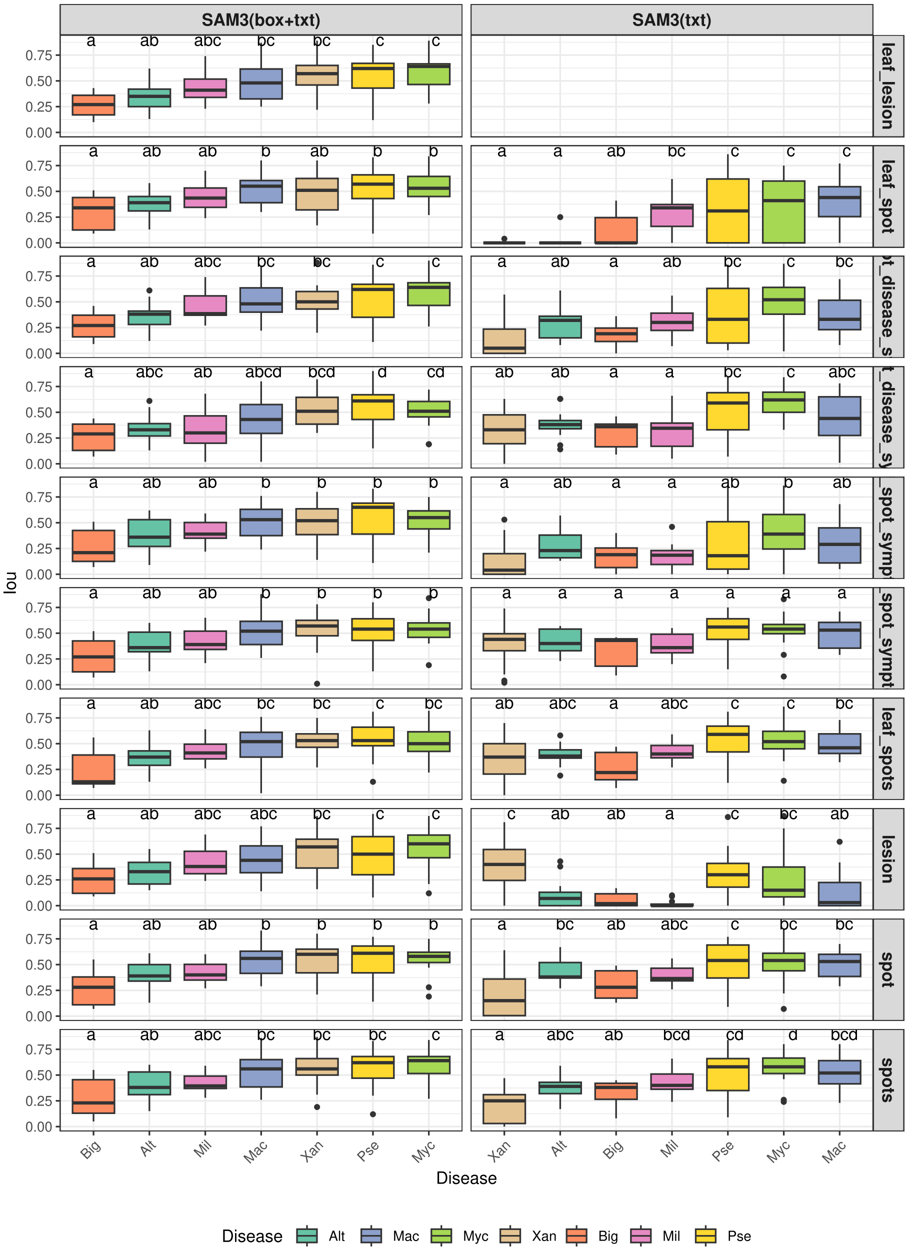


Figure 6: Segmentation performance (IoU) of the 19 Model_Prompt combinations (10 SAM3(box+txt) and 9 SAM3(txt)) per oilseed rape leaf disease. Boxplots show the distribution of IoU values, letters above boxes indicate homogeneous groups with no significant difference from post-hoc pairwise comparisons.

### D Complements on the estimation of diseased surface experiment

#### D.1 Generalization experiment

To assess the generalization of the pseudo-mask generation strategies in terms of diseased surface estimation, we applied the same procedure used for the oilseed rape dataset (detailed in Section ??) to the apple dataset. The resulting rRMSE values are presented in Table 5.

Table 5: Evaluation of the estimation of the diseased surface on the apple dataset using rRMSE as the metric.

| Model | rRMSE |
|---|---|
| FS | 0.2103 |
| CAM + SAM(box) | 1.0615 |
| CAM + SAM3(box) | 1.0849 |
| SAM3(txt)_best_prompt | 0.6244 |
| CAM + SAM3(box + txt)_best_prompt | 0.7219 |

The fully supervised model again achieved the best performance, serving as the baseline, with an rRMSE of 0.2103. The CAM-based method alone achieved performance similar to that observed on the oilseed rape dataset. However, contrary to the oilseed rape dataset, the text-only model (SAM3(txt)) outperformed the hybrid approaches on the apple dataset. This result is consistent with the segmentation performance (IoU) observed on this dataset.

#### D.2 Estimation of the bias

To assess whether a model tends to over- or under-estimate the symptom surface, a linear regression is fitted between the predicted ratio $\rho_{\text{pred}}$ and the reference ratio $\rho_{\text{ref}}$ across all images of the dataset:

$$\rho_{\text{pred}} = \beta \cdot \rho_{\text{ref}} + \mu \tag{3}$$

where $\beta$ is the slope and $\mu$ is the intercept. A slope $\beta > 1$ indicates a tendency to over-estimate the symptom surface relative to the manual annotation, while $\beta < 1$ indicates under-estimation. A non-zero intercept $\mu$ reveals a constant offset between the predicted and reference ratios. A model with no systematic bias would ideally satisfy $\beta = 1$ and $\mu = 0$.

This linear regression is fitted for each segmentation model trained with the different batches of pseudo-masks, as well as for the fully supervised baseline.

**Results**

Figure 7 presents the linear regression for the fully supervised model, the CAM-based-only model, and the best-prompt models for the text-guided approaches (strict and hybrid). Figures 8 and 9 present the linear regressions for all prompts of the CAM + SAM3(txt) and CAM + SAM3(box+txt) models, respectively.

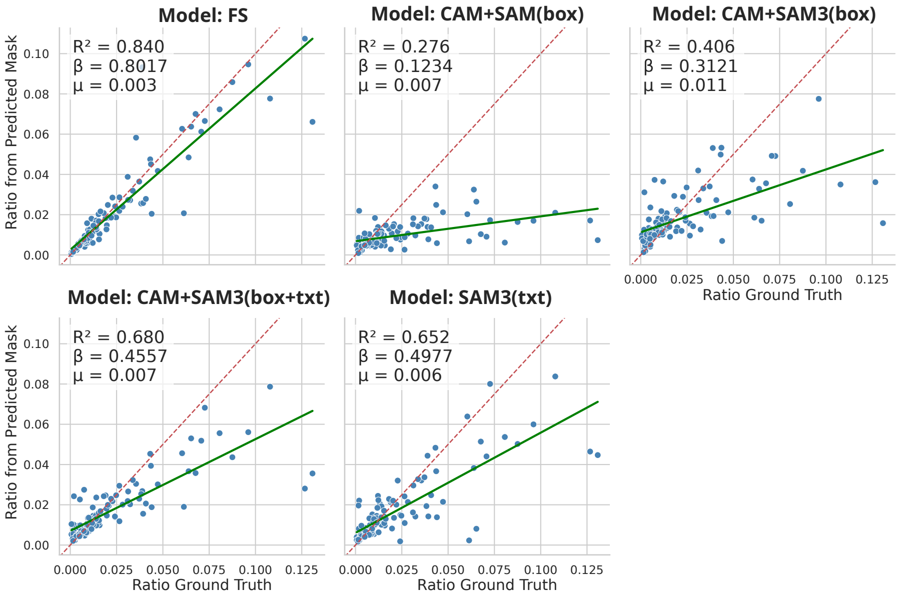


Figure 7: Linear regression between predicted and reference diseased-surface ratios for the fully supervised model, the CAM-based-only model, and the best-prompt text-guided models (strict and hybrid).

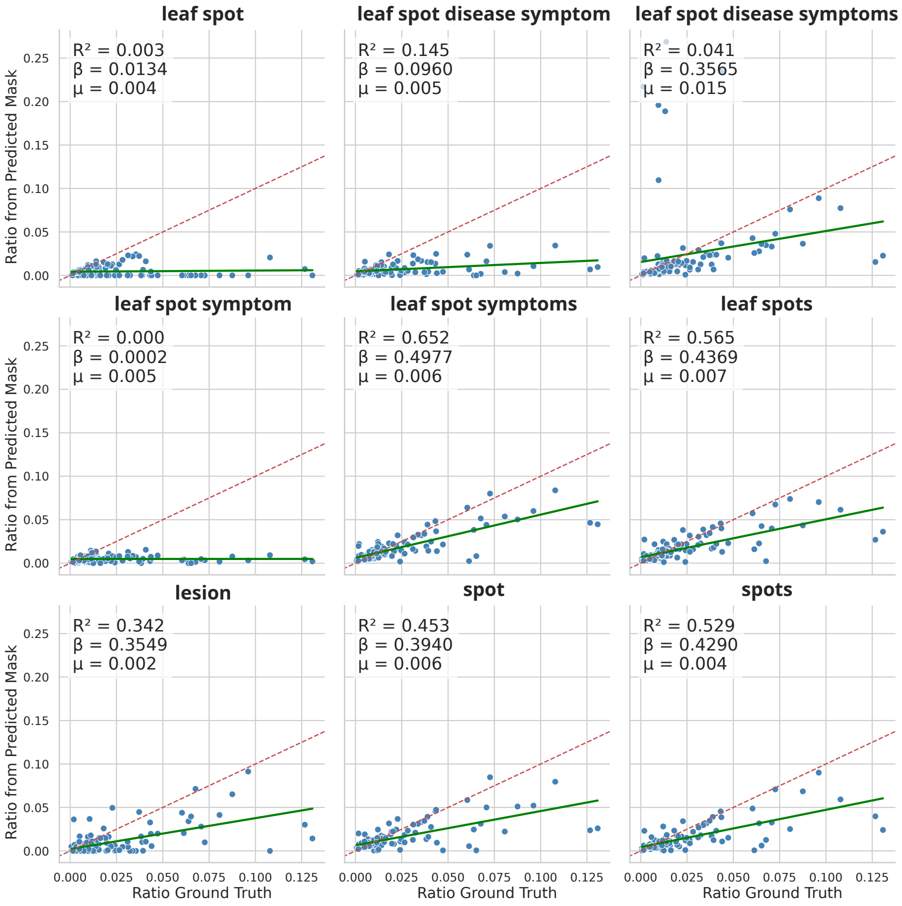


Figure 8: Linear regression between predicted and reference diseased-surface ratios for all text prompts of the CAM + SAM3(txt) models.

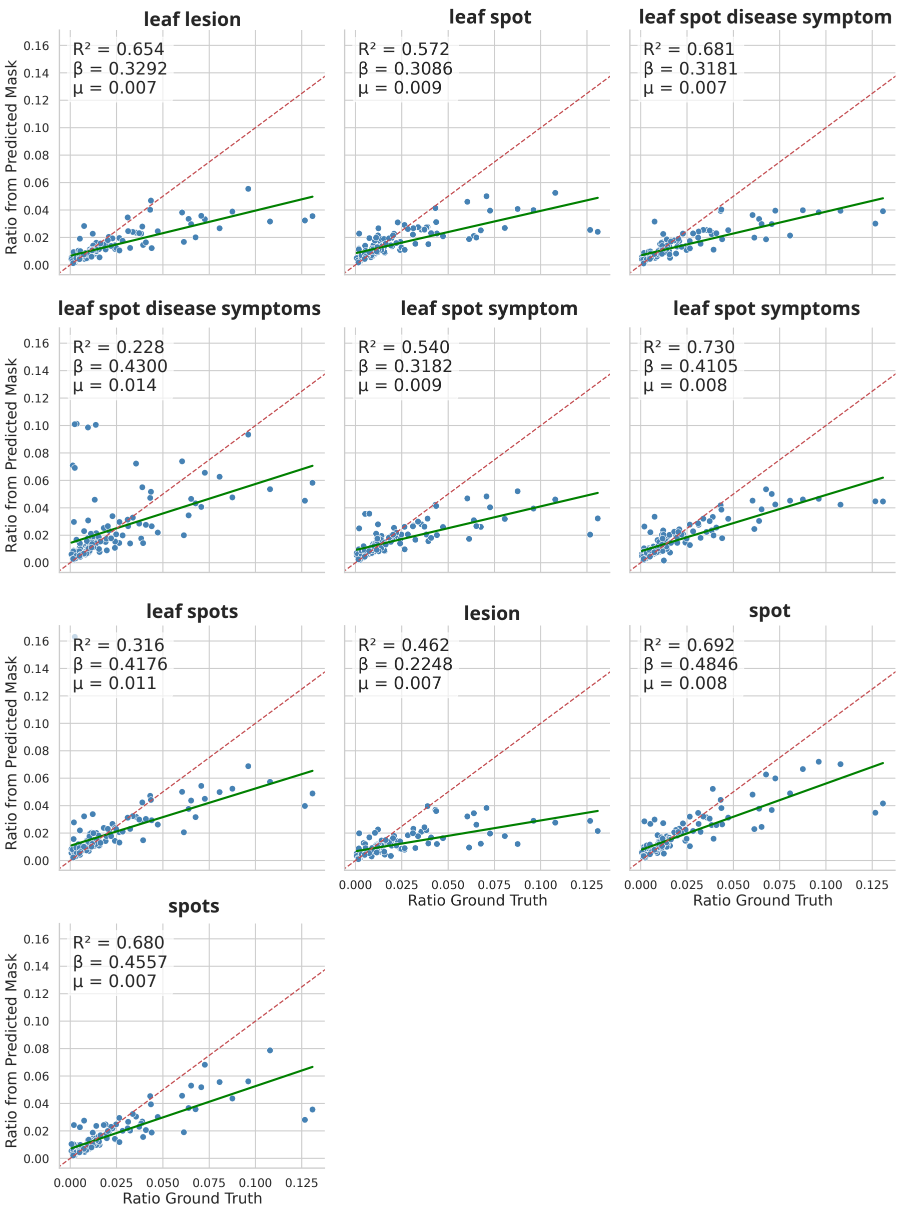


Figure 9: Linear regression between predicted and reference diseased-surface ratios for all prompts of the CAM + SAM3(box+txt) models.
